# A human neuron-microglia tri-culture platform to study the influence of microglia on developing neuronal networks *in vitro*

**DOI:** 10.64898/2026.04.06.716675

**Authors:** Sara Guerrisi, Adam Pavlinek, Olivia L. Cunningham, George Chennell, Anthony C. Vernon, Deepak P. Srivastava

## Abstract

Human brain function is dependent on synaptic architecture and function between a range of different cell types. Glutamatergic and GABAergic neurons provide the basis by which the excitatory and inhibitory balance is achieved in cortical networks, and microglia interact with them to shape synaptic architecture and neural networks. Understanding the interactions between these cell types is crucial to elucidating mechanisms relevant to brain physiology and, potentially, to neurodevelopmental and neurological disorders. Here, we establish a rapid and reproducible human tri-culture platform comprising deterministically-programmed glutamatergic neurons, GABAergic neurons, and microglia to facilitate cell-cell interaction studies during human cortical development. Using these deterministically-programmed ioCells, we systematically optimised neuronal ratios, culture conditions, and the timing of microglial integration to generate a stable neuronal network prior to microglia incorporation. Multi-electrode arrays (MEAs) recordings identified an 80:20 glutamatergic-to-GABAergic ratio as the most robust configuration for sustained and reproducible network activity in this context. Structural characterisation using automated high-content imaging confirmed the formation of both excitatory and inhibitory synapses, while longitudinal MEA recordings demonstrated stable network maturation following microglial incorporation. Microglia incorporation influenced neuronal firing dynamics, increasing burst activity without disrupting early synapse formation. As a proof of concept for disease modelling, we incorporated microglia carrying the Alzheimer’s disease-associated *TREM2* R47H mutation and detected subtle but reproducible alterations in neuronal burst dynamics. Together, this work establishes a defined human neuron-microglia triculture platform that enables scalable investigation of neuroimmune interactions and genetic variants, laying the foundations for more complex future models.

## Introduction

Microglia are brain resident innate immune cells (Matuleviciute *et al*., 2023). They represent 5-10% of all the cells in the central nervous system (CNS), and 10-15% of all glial cells (Nayak *et al*., 2014; Mondelli *et al*., 2017). Microglia are suggested to play essential roles in brain development, homeostasis, and responses to environmental perturbation (Nayak *et al*., 2014; Hanger *et al*., 2020; Upthegrove *et al*., 2025). Beyond immune surveillance, microglia actively regulate synaptic formation, elimination, and remodelling, contributing to circuit refinement during development and influencing neuronal function (Paolicelli *et al*., 2011; Wohleb, 2016; Eyo & Molofsky, 2023). Dysregulation of microglia–synapse interactions has been implicated in a wide range of neurodevelopmental and neuropsychiatric disorders, including schizophrenia, autism spectrum disorders, and neurodegenerative diseases (Sellgren *et al*., 2019; Hanger *et al*., 2020; Couch *et al*., 2024; Teter *et al*., 2025; Upthegrove *et al*., 2025; Zhong *et al*., 2025). In particular, there is increasing evidence that microglia actively sense and respond to changes in neuronal activity but also have the potential to influence neural activity following a change in their own functional state (Badimon *et al*., 2020; Que *et al*., 2024). However, our understanding of the role of microglia in these processes during human neurodevelopment remains limited (Matuleviciute *et al*., 2023; Hume, 2025; Rittenhouse *et al*., 2025). Specifically, whilst significant advances in understanding neuron-microglia interactions have been made (Kuhn *et al*., 2004; Pascual *et al*., 2012; Schafer *et al*., 2012), many of these mechanistic insights have come from rodent models. These are limited by species-specific differences in immune signalling, synaptic architecture, and developmental timing (Breschi *et al*., 2017; Deng, 2017; Beninger *et al*., 2024; Cserép *et al*., 2025). Consequently, there is increasing demand for reproducible, human-derived *in vitro* model systems that enable controlled investigation of neuron–microglia interactions in the context of human neurodevelopment (Hanger *et al*., 2020; Hasselmann & Blurton-Jones, 2020; Couch *et al*., 2024).

To address this gap, human induced pluripotent stem cell (hiPSC)-based models are used extensively to study neuronal and glial function and interactions in the context of physiology, as well as disease modelling and drug discovery (Faust *et al*., 2025; Li *et al*., 2025; Lish *et al*., 2025b; Tegtmeyer *et al*., 2025). Numerous protocols now exist for differentiating excitatory glutamatergic neurons, inhibitory GABAergic neurons, astrocytes, and microglia from human pluripotent stem cells (Odawara *et al*., 2014; Abud *et al*., 2017; Hasselmann & Blurton-Jones, 2020; Mossink *et al*., 2022). Forward programming approaches in particular have facilitated rapid and scalable production of specific neuronal and glial cell populations (Pawlowski *et al*., 2017; Mossink *et al*., 2021; Cerina *et al*., 2023; Habich *et al*., 2025), supporting functional studies of synapse function and dysfunction (Sellgren *et al*., 2019; Berryer *et al*., 2023), significantly reducing culture time and variability (Mordelt *et al*., 2025). In combination with high content imaging approaches (Nieland *et al*., 2014; Papandreou *et al*., 2023; Tegtmeyer *et al*., 2025), as well as multi-electrode arrays (MEAs) (Pavlinek *et al*., 2026), it is now possible to carry out detailed functional studies on cultures with multiple neural cell types (Lish *et al*., 2025b; Mordelt *et al*., 2026; Wu *et al*., 2026).

Recent work in the field has highlighted the importance of establishing appropriate excitation/inhibition (E/I) balance in neuronal cultures (Yuan *et al*., 2023; Pavlinek *et al*., 2026). This has demonstrated that inclusion of inhibitory neurons significantly alters network maturation, burst dynamics, and pharmacological responsiveness (Mossink *et al*., 2021; Cerina *et al*., 2023). In parallel, human tri-culture systems incorporating neurons, astrocytes, and microglia have been developed to model neuroinflammatory signalling and glial crosstalk (Luchena *et al*., 2022; Lish *et al*., 2025b; Zheng *et al*., 2025). These studies underscore the value of increasing cellular complexity *in vitro* to better approximate human brain physiology. However, there is a need for standardised, scalable, and experimentally tractable human models specifically designed to interrogate neuroimmune interactions at the level of defined neuronal networks (Summers *et al*., 2024; Haskell *et al*., 2026). Many existing co-culture systems either rely on long differentiation timelines, include rodent astrocytes as additional regulatory elements, or use excitatory neuronal monocultures without systematic control of inhibitory composition (Li *et al*., 2025; Lish *et al*., 2025b). A fast and standardised E/I-balanced neuronal system integrated with human microglia represents a practical and complementary approach for focused neuroimmune studies, enabling investigations of the effects of microglia’s on neuronal activity, both physiologically, and in response to perturbations.

Here, we present a rapid, reproducible, and scalable system comprising deterministically-programmed hiPSC-derived glutamatergic neurons, GABAergic neurons, and microglia (“ioCells”, bit.bio). We systematically optimised neuronal ratios, culture conditions, and the timing of microglial integration to generate a stable and functionally active neuronal network prior to immune cell incorporation. MEA-based screening of glutamatergic/GABAergic ratios identified an 80:20 configuration as the most robust for sustained and reproducible activity, consistent with physiological inhibitory neuron proportions in the developing human cortex (Marin, 2025). Microglia were subsequently integrated into this defined E/I-balanced co-culture, enabling controlled assessment of their effects on synaptogenesis, electrophysiological maturation, and emergent network dynamics. To support standardised structural analysis, we further developed an automated high-content imaging pipeline (Couch *et al*., 2024; Sichlinger *et al*., 2024) to quantify excitatory and inhibitory synapses and to characterise microglial morphology. In this tri-culture system, we observed the formation of both excitatory and inhibitory synapses, with structural stability maintained throughout differentiation. Furthermore, this system exhibited sustained network activity with reproducible modulation of burst properties following microglial incorporation. To demonstrate the application of this system in a disease context, we further incorporated *TREM2* R47H mutant microglia to examine how genetic perturbation of microglia impacted synaptic and network properties. This protocol provides a fast, tractable and high-throughput-compatible system for studying human neuron-glia interactions relevant to neurodevelopmental and neurodegenerative processes, aligning with the growing adoption of New Approach Methodologies (NAMs).

## Material and Methods

### Triculture Protocol

#### Neuronal culture

ioGlutamatergic neurons (bit.bio, io1001) and ioGABAergic neurons (bit.bio, io1003) were thawed and seeded onto plates pre-coated with Poly-D-Lysine (Gibco, A3890401) for 3 hours at 37°C, washed with deionised water and dried. A second layer of coating was added with Geltrex (Gibco, A1413302) diluted in DMEM F/12 (Sigma-Aldrich, D6421-500ML) 1:100 for 1 hour at 37°C. Cells were directly thawed and cultured on PhenoPlate 96 well (Revvity, 6055300) for high-content imaging, or CytoView MEAs 24 well Plate (Axion) for MEA recording.

For monoculture of ioGlutamatergic neurons, a total of 35,000 cells were seeded per well for the PhenoPlate 96, and 150.000 cells were seeded per well of CytoView MEAs 24 well Plate. Cells were resuspended in CompGN+D+Rocki media, composed of Neurobasal (Gibco, 21103049), Glutamax (100X, Gibco, 35050061), 2-Mercaptoethanol (Millipore Sigma, M3148, 25 μM), B27 (50X, Gibco, 17504044), NT3 (10ng/mL) BDNF (Biolegend, 788902, 5ng/mL), Doxycyline, and Y-27632 (10 μM). In CytoView MEAs 24 well plate cells were seeded using the drop method, with a 50 µL drop of cells incubated at 37°C overnight.

For monoculture of ioGABAergic neurons, 48,000 cells were seeded in each well of a PhenoPlate 96 well, and 150,000 cells per well were seeded on CytoView MEAs 24 well plates. Cells were resuspended with CompGS+D+Rocki media composed of DMEM/F12 (Sigma-Aldrich, D6421-500ML), N-2 Supplement (Gibco, 17502048, 100X), MEM-NEAA (100X, ThermoFisher, 11140-035), Doxycyline and Y-27632 (10 μM). The day after, 500 µL per well of CompGS+D without Rocki was added to the CytoView MEA plate, and 200 µL per well was used for a full media change per well of the PhenoPlate 96. From day 3 and onwards, a half media change was performed three times a week with CompMM media composed of BrainPhys, B27 (50X), Glutamax (100X) and BDNF.

To co-culture ioGlutamatergic neurons with ioGABAergic neurons, 35.000 and 7.000 cells respectively were added to the PhenoPlate 96, and 150.000 and 30.000 cells respectively were seeded on CytoView MEA on day 0. The day after, 500 µL per well of CompGN+D without Rocki was added to the CytoView MEA plate, and 200 µL per well was used for full media change on PhenoPlate 96 well. On day 2, a full media change was performed with CompGN+D with DAPT (Thermo Scientific Chemicals, J65864.MA). From Day 3 and onwards, half-media change was performed three times a week with CompGN(BrainPhys), composed of BrainPhys (Stemcell Technologies, 05790), Glutamax (100X), 2- Mercaptoethanol, B27 (50X), NT3 and BDNF.

#### Microglia culture

ioMicroglia (bitbio, io1021) were received at King’s College London, thawed and seeded onto plates pre-coated with Poly-D-Lysine (Gibco, A3890401) for 3 hours at 37°C, washed with deionised water and dried, followed by a second layer of coating with Geltrex (Gibco, A1413302) for 1 hour at 37°C. A total of 375,250 cells per well were seeded per well of a 6-well plate and resuspended in media composed of Advanced-DMEM/F12 (Gibco, 12491015), Glutamax (Gibco, 35050061), N-2 Supplement (100X), 2-Mercaptoethanol, B27 (50X), Recombinant Human M-CSF (Peprotech, 300-25, 50 ng/mL), Doxycycline, and Y-27632 (10 μM). The day after, a full media change was performed with CompMicroglia media comprised of Advanced DMEM supplemented with Glutamax (100X) and N-2, 2-Mercaptoethanol, M-CSF (10 ng/mL), and Recombinant Human IL-34 (Peprotech, 200-34, 100 ng/mL). On day 3, ioMicroglia were ready to be incorporated in the Triculture model. For longer-term ioMicroglia monoculture, we performed a half-media change with CompMicroglia every 72h supplemented with M-CSF (20 ng/mL), and IL-34 (200 ng/mL).

#### Tri-culture platform of ioGlutamergic, ioGABAergic neurons and ioMicroglia

After 10 days of neuronal co-culture, ioMicroglia were added to the model. In brief, ioMicroglia in monoculture were detached using Accutase for 4 minutes at 37°C and resuspended in DMEM. After centrifugation at 1200 rpm for 5 minutes, the cell pellet was resuspended with Tric(ADMEM) media composed of Advanced DMEM/F12, Glutamax, N-2 Supplement, B27, 2-Mercaptoethanol, NT3, BDNF, IL-34 and M-CSF, and cells were seeded at a ratio of 1:5 microglia to neurons. After 72h a full media change was performed with Tric(BrainPhys) composed of BrainPhys, Glutamax, 2-Mercaptoethanol, B27, NT3, BDNF, IL-34 and M-CSF. From day 16 and onwards, half media change was performed three times a week.

Full details concerning the culturing protocol may be found on protocols.io at https://www.protocols.io/edit/protocol-triculture-model-bit-bio-for-96-well-plat-hb66b2rhf

### High-content imaging

#### Immunocytochemistry

Cells were fixed with a solution containing 1X PBS with Ca2+ and Mg+, 4% PFA and 4% Sucrose for 9 minutes at room temperature, following 2X washes with PBS with Ca2+ and Mg+, and stored at 4°C. Prior to incubation with primary antibodies, cells were incubated in a blocking solution with 0.1% Triton-X, 4% Normal Goat or Donkey Serum, and 1X PBS, for 1 hour at room temperature. Cells were then incubated overnight with diluted primary antibodies at 4°C (Supplementary Table 1). Cells were then washed 3X (5 minutes) with 1X PBS with Ca2+ and Mg+, and incubated with secondary antibodies diluted 1:750 for 1 hour at room temperature. Three further washes with 1X PBS with Ca2+ and Mg+ were performed, followed by a 5-minute incubation with PBS+DAPI (1:20,000). Finally, cells were washed twice and kept in PBS for imaging. Details about primary and secondary antibodies and their dilution used can be found in **Supplementary Table 1**. Each experiment and combination of antibodies was performed with a No Antibody control, No Primary antibody and No Secondary antibody control.

#### High-content microscopy

Immunostained cells were imaged using the OperaPhenix High-Content Screening System (Revvity). For each staining condition, 3 wells were prepared and treated as technical replicates. Cells were imaged with either a 20x Air objective (NA 0.4), 40x water objective (NA 1.1), or a 63x water objective (NA 1.15), with between 15 and 30 fields of view per well. Laser power and exposure time were adjusted to ensure an optimal signal-to-noise ratio and kept consistent across the plate.

#### Image Analysis

Image analysis was performed using Harmony High-Content Imaging and Analysis Software. A custom analysis pipeline was developed using representative images randomly selected from all available fields of view and verified against control wells. The pipeline was then run automatically across all the images, to ensure robustness of results. For synaptic quantification analysis, nuclei were found using DAPI channel, filtering out the DAPI+ nuclei on the border of the image. MAP2+ staining was used to find the cell body and neurites of neurons, in order to define our ROI. Consequently, puncta were detected within the ROI using the pre-synaptic protein bassoon and Vesicular inhibitory amino acid transporter (VGAT) and post-synaptic NMDA receptor subunit GluN1 (excitatory synapses) and neuroligin 2 (Nlgn2 - inhibitory synapse) using either the 488nm or 568nm channel. In each channel, spots were filtered by size and their intensity defined as Spot to Background signal. A size filter was set between 27 and 100 pixels, equivalent to 0.8-5 µm^2^. The intensity threshold was set according to that measured from the No Secondary Antibody control well. Selected puncta were then masked on top of each other, in order to count co-localisation, defined as >50% overlap (Couch *et al*., 2024; Sichlinger *et al*., 2024). Morphology features were extracted from MAP2+ dendrite staining to calculate the density of synapses defined as number of overlapping puncta / average neurite length as previously reported elsewhere (Nieland *et al*., 2014; Couch *et al*., 2024). For microglia morphology analysis, DAPI+ nuclei were identified, filtering out those on the border of the image. Then the IBA1 channel was used to localise microglia, and the Find Neurites block was used to map and measure microglia’s branches.

### Live-cell calcium imaging

To perform live-cell calcium imaging we used the Fluo-4 AM dye (Invitrogen, F14201). Cells were seeded onto Ibidi µ-Slide 8 Well culture slides (Ibidi, 80806). On DIV25 of neuronal culture, one vial of Fluo-4 AM was resuspended in 46 µl of DMSO to generate a 1mM stock. The stock was diluted 1:200 in culture media and incubated at 37°C for 1h in the dark. Then, a full media change was been performed twice, every 15 minutes, with cells left to equilibrate for 30 minutes in the incubator prior to imaging. All Imaging was performed with a Ti-2 Three Camera Calcium Microscope from Nikon using a 40x water objective.

### Multi Electrode Arrays (MEAs)

Cells were seeded onto CytoView MEAs 24 well Plate as described above. MEA recordings were obtained across all electrodes using the Axion Maestro and Axis Navigator software (12.5 kHz sampling rate, 200-3000 Hz band-pass). The Axion Maestro was stabilised at 37°C and 5% CO2 prior to recording, and recordings were always made prior to any media change. For comparison of spontaneous activity between different culture conditions, daily 10 minute recordings were obtained between DIV14 to DIV27. Recordings consisted of N=4 independent replicates derived from independent differentiations, each of them with 4 different wells to account for technical variability (16 wells total).

The recording files were then analysed in batch with Axis Navigator Software for spontaneous electrical activity and burst detection (Pavlinek *et al*., 2026). The resulting .csv files were used to run statistical analysis and plots.

### Statistics

Results were analysed from N=3 or N=4 independent replicates for each experiment, defined as independent differentiations from different vials of ioCells. For high-content imaging results, data from the individuals fields of view within each technical replicate (well) were averaged and subsequently normalised to a baseline control, specifically either ioGlutamatergic only or ioGABAergic only conditions. Dependent variables were compared across culture conditions using either a one sample t-test or a non-parametric Kurskal-Wallis test test as appropriate in GraphPad Prism 9 for Windows (GraphPad Software LLC, California, USA), with α = 0.05.

Statistical analysis of MEA recording data was performed using R version 4.4.2, with an in-house custom script developed using R Studio. For each parameter of interest, the average of technical replicates (4-6 wells) was calculated, and the N=4 biological replicates (independent differentiations from different vials of ioCells) were plotted in a line graph across multiple time points. The MEA data was not normally distributed (Shapiro-Wilk Test p<0.05), hence we performed a non-parametric Aligned Rank Transformation (ART) test, using the *ARTool* R package with α = 0.05. Where appropriate, *post-hoc* Pairwise Contrasts were performed across conditions for each timepoint. For time-specific comparison, R was used to build boxplots using the *ggplot* package. Details of which test is performed are detailed in legend figures. Details of statistical analysis for figures 2,3, 4 and 5 can be found in **Supplementary Tables 2-6**.

## Results

### Integrating microglia into a two-dimensional culture of glutamatergic and GABAergic neurons

Prior to generating the tri-culture platform, we optimised the neuronal co-culture without microglia. First, we tested different ratios of ioGlutamatergic and ioGABAergic neurons, to determine the most efficient ratio of these cell types that promotes a functional neural culture with both excitatory and inhibitory components. Immediately after thawing, ioGlutamatergic and ioGABAergic neurons (bit.bio) were co-seeded onto multi-electrode array (MEA) culture plates, at the following ratios: 80:20, 70:30, 60:40 and 50:50 (**Figure 1A**). Co-cultures were subsequently monitored longitudinally for network activity, and mean firing rate (MFR), burst frequency, and synchrony index were quantified (Pavlinek *et al*., 2026). Across independent biological replicates (N=4), we observed that the 80:20 configuration consistently produced the most robust activity profile, characterised by stable active electrode numbers and sustained burst activity over time (**Figure 1B; Supplementary Figure 2**). Unsurprisingly, increasing the proportion of inhibitory neurons resulted in reductions in overall firing rates and decreases in burst stability (Figure 1B). In contrast, reducing the proportion of inhibitory neurons led to increased variability in network activity (**Figure 1B; Supplementary Figure 2**). Based on these data, we selected the 80:20 ratio for all subsequent triculture experiments as it provided a physiologically relevant (Mordelt *et al*., 2026) and functionally stable E/I-balanced neuronal co-culture (Cerina *et al*., 2023)

**Figure 1.**
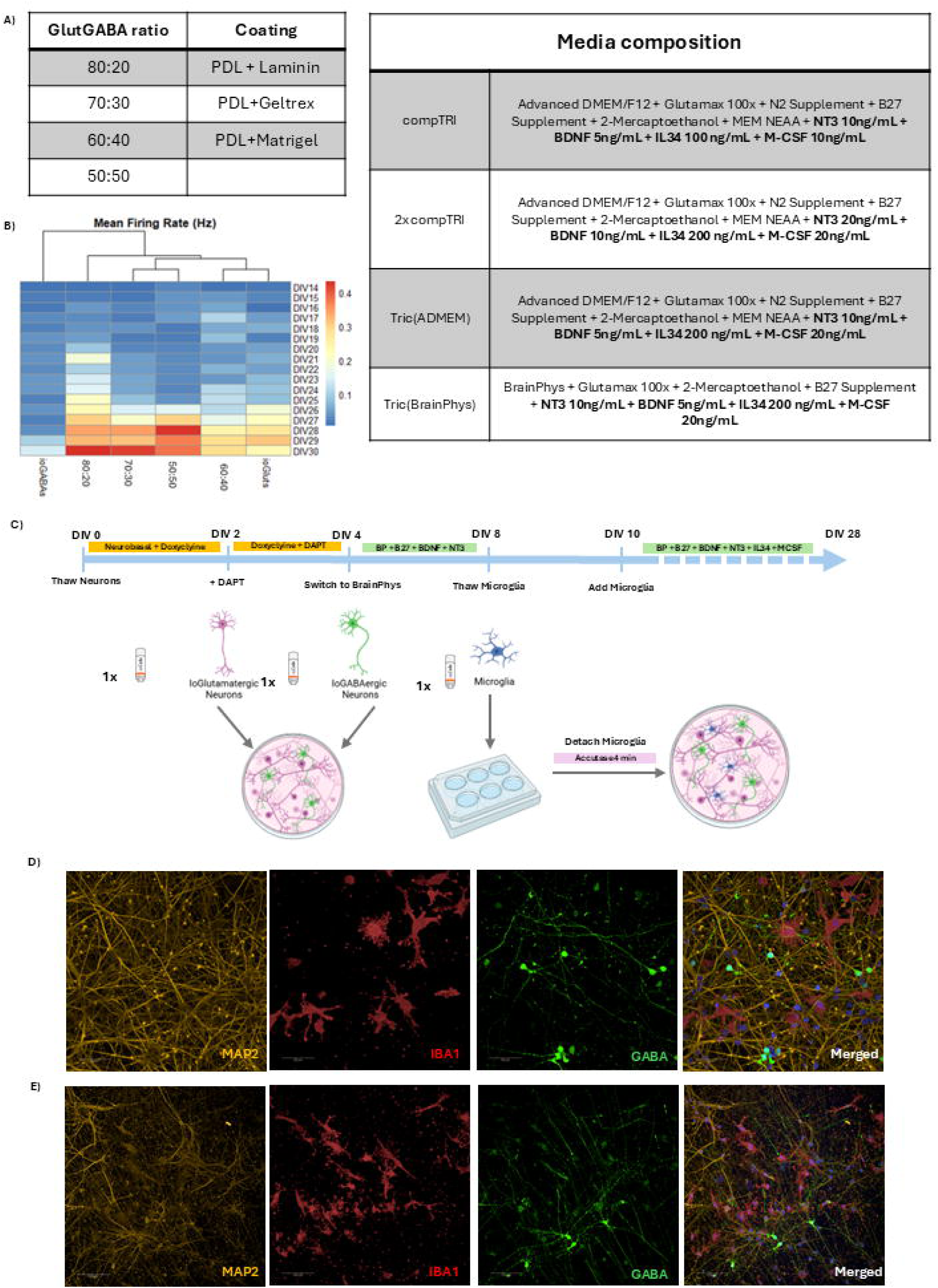
Optimisation of protocol to integrate microglia into a co-culture of ioGlutamatergic and ioGABAergic neurons. **A)** Table summarising conditions tested for cell ratio, coating and media composition. **B)** Heatmap showing mean firing rate (MFR) of ioGlutamatergic (ioGlut), ioGABAergic (ioGABA) neurons and varying rations of ioGlut and ioGABA co-cultures. **C)** Schematic of the tri-culture protocol. Figure shows media composition, timeline and seeding approach. **D)** Representative images of Tri-culture at DIV21 stained for MAP2 (orange, neuron), IBA1 (red, microglia) and GABA (green, GABAergic neuron), and merged. Imaged with 40x water lens. **E)** Representative images of Tri-culture at DIV28, 20x Air lens.

Next, we optimised the culture conditions to support simultaneous maintenance of neurons and microglia (**Figure 1A**). Initial experiments used Advanced DMEM/F12-based triculture media (compTRI) with double concentrations of NT3, BDNF, IL34 and M-CSF to account for the half media change (**Figure 1B**). However, it was observed that these high concentrations of NT3 and BDNF led to formation of significant clusters of neurons (**Supplementary Figure 1)**. These substantially decreased when only IL-34 and M-CSF concentrations were doubled in the CompTRI media (Tric[ADMEM]). Despite this, activity in the tri-cultures cultured in Advanced DMEM/F12-based media was suboptimal when checked with live-cell Ca2+ imaging (**Supplementary Video 1**). Substitution of Advanced DMEM/F12 with BrainPhys medium restored spontaneous firing and sustained bursting behaviour (**Figure 1B),** resulting in a finalised triculture medium (Tric[BrainPhys]) compatible with both neuronal activity and microglial viability (**Figure 1C**).

Using the optimised conditions described above, ioGlutamatergic and ioGABAergic neurons were co-seeded at DIV0 at a 5:1 ratio (80:20) (Crocco *et al*., 2025; Mordelt *et al*., 2026). ioMicroglia were thawed at DIV8, expanded for 48 hours and incorporated into neuronal cultures at DIV10 (**Figure 1C**). This staged integration ensured the establishment of a stable neuronal network prior to the addition of microglia. Immunocytochemical analysis at DIV21 confirmed the presence of MAP2-positive neuronal networks, GABA-positive inhibitory neurons, and IBA1-positive microglia (**Figure 1D**). ioMicroglia were observed to be evenly distributed across the culture surface and displayed a characteristic ramified morphology (Figure 1D). Importantly, the tri-culture remained structurally stable through DIV28, which represent a mature network timepoint for ioGlutamatergic neurons with persistent integration of microglia and maintained neuronal network architecture (**Figure 1E**).

### Triculture model correctly forms both excitatory and inhibitory synapses

To determine whether including microglia affected synaptogenesis, we performed a systematic structural characterisation of synapses across multiple culture conditions. Mono-, co- and tri-cutlures of ioGlutamertgic neurons, ioGABAergic neurons and ioMicroglia were seeded in parallel and analysed at DIV21 using high-content imaging (**Figure 2**). Excitatory synapses were identified by assessment of overlapping bassoon (pre-synaptic) and the NDMA receptor subunit GluN1 (post-synaptic). Inhibitory synapses were identified by overlapping VGAT (pre-synaptic) and Nlgn2 (post-synaptic) (**Figure 2A**). Building on previously published synapse detection algorithms (Nieland *et al*., 2014; Couch *et al*., 2024; Sichlinger *et al*., 2024) we developed an automatic imaging pipeline to quantify the linear density of pre-, post-synaptic and colocalised (synaptic) puncta (**Figure 2B**). Images were segmented to detect neurites and surrounding regions, and synaptic puncta were quantified when pre- and post-synaptic puncta overlapped by more than 50%.

**Figure 2.**
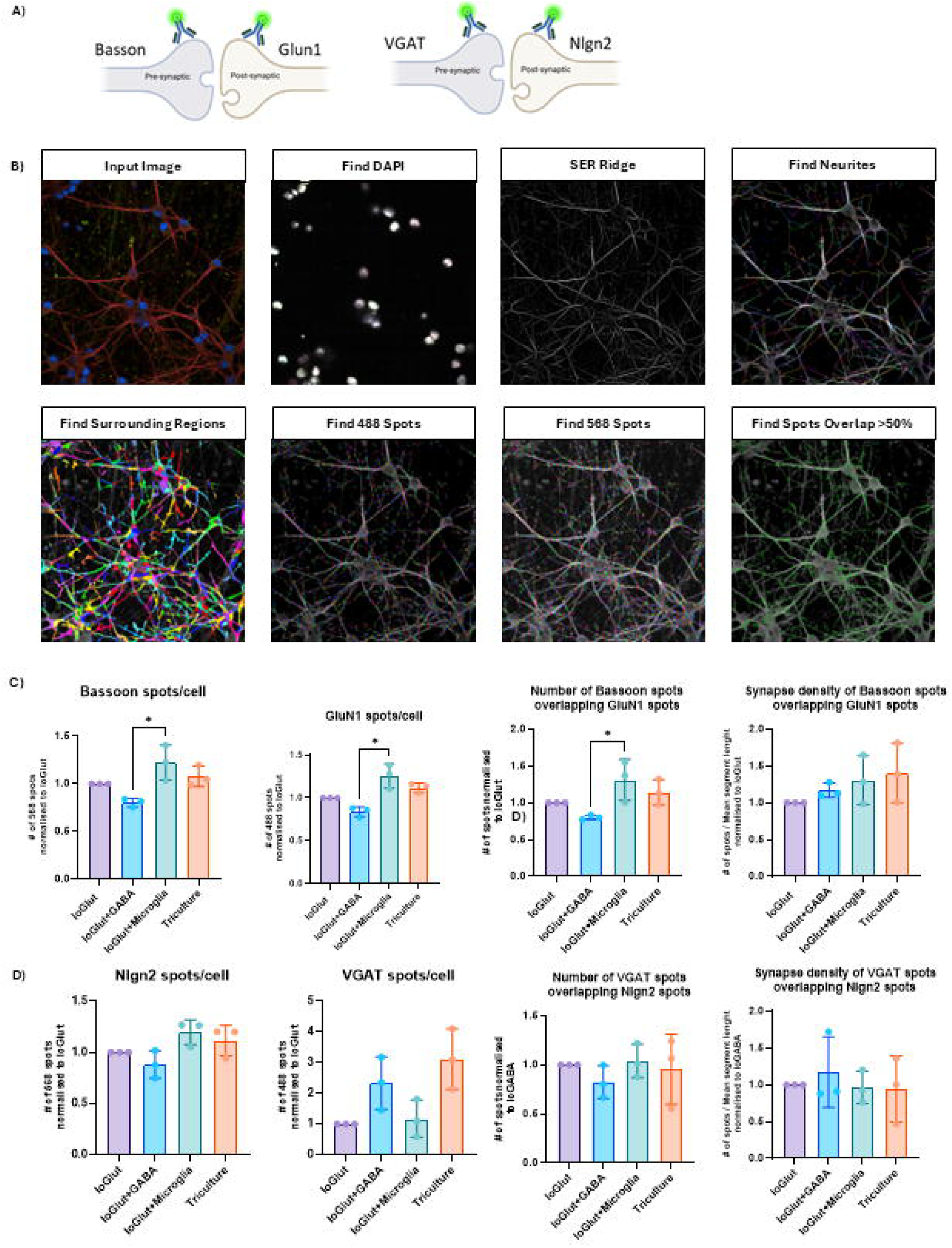
Tri-culture model correctly forms both excitatory and inhibitory synapses. **A)** Schematic representation of the pre- and post-synaptic proteins used to mark excitatory and inhibitory synapses in this experiment. **B)** Overview of the imaging analysis pipeline developed to analyse puncta co-localisation in the orange and green channel, in MAP2+ dendrites (red). **C)** Results of excitatory and **D)** inhibitory synapses across different culture conditions of neurons in the presence and absence of microglia, normalised to ioGlutamatergic (ioGlut) only condition (Kruskal-Wallis test, with Dunn’s post hoc test, N=3). Single protein spot are plotted normalised per number of cells, and as Fold Change to ioGlut condition, number of co-localised puncta and synapse density are normalised to ioGlut condition.

Using this pipeline, we analysed data from N=3 wells per condition, with N=30 fields of view per well, from N=3 independent biological replicates. Quantification revealed a statistically significant difference in the abundance of excitatory synaptic proteins between ioGlutamatergic+ioGABAergic compared to ioGlutamatergic+ioMicroglia conditions. (**Figure 2C**). Consistent with this, the number of Bassoon puncta colocalising with GluN1 was also significantly elevated, suggesting an increased formation of purported excitatory synapses. However, when normalised to neurite length, overall excitatory synapse density did not significantly differ across culture conditions (**Figure 2C**). Incorporation of microglia into ioGlutamatergic+ioGABAergic co-cultures did not significantly alter excitatory synaptic protein expression, with the tri-culture displaying similar levels of synaptic protein and synapse density to co-cultures.

We next examined the density of inhibitory synaptic protein and synapse expression across all culture conditions, defined as total number of overlapping puncta divided by the average dendrite length (Nieland *et al*., 2014; Couch *et al*., 2024; Sichlinger *et al*., 2024). This revealed that the expression of VGAT and Nnlgn2 was broadly comparable across all culture conditions (**Figure 3A & B**). Similarly, the density of synapses, assessed as the number of VGAT puncta colocalising with Nlgn2 did not differ across culture conditions (**Figure 3A & B**).

**Figure 3.**
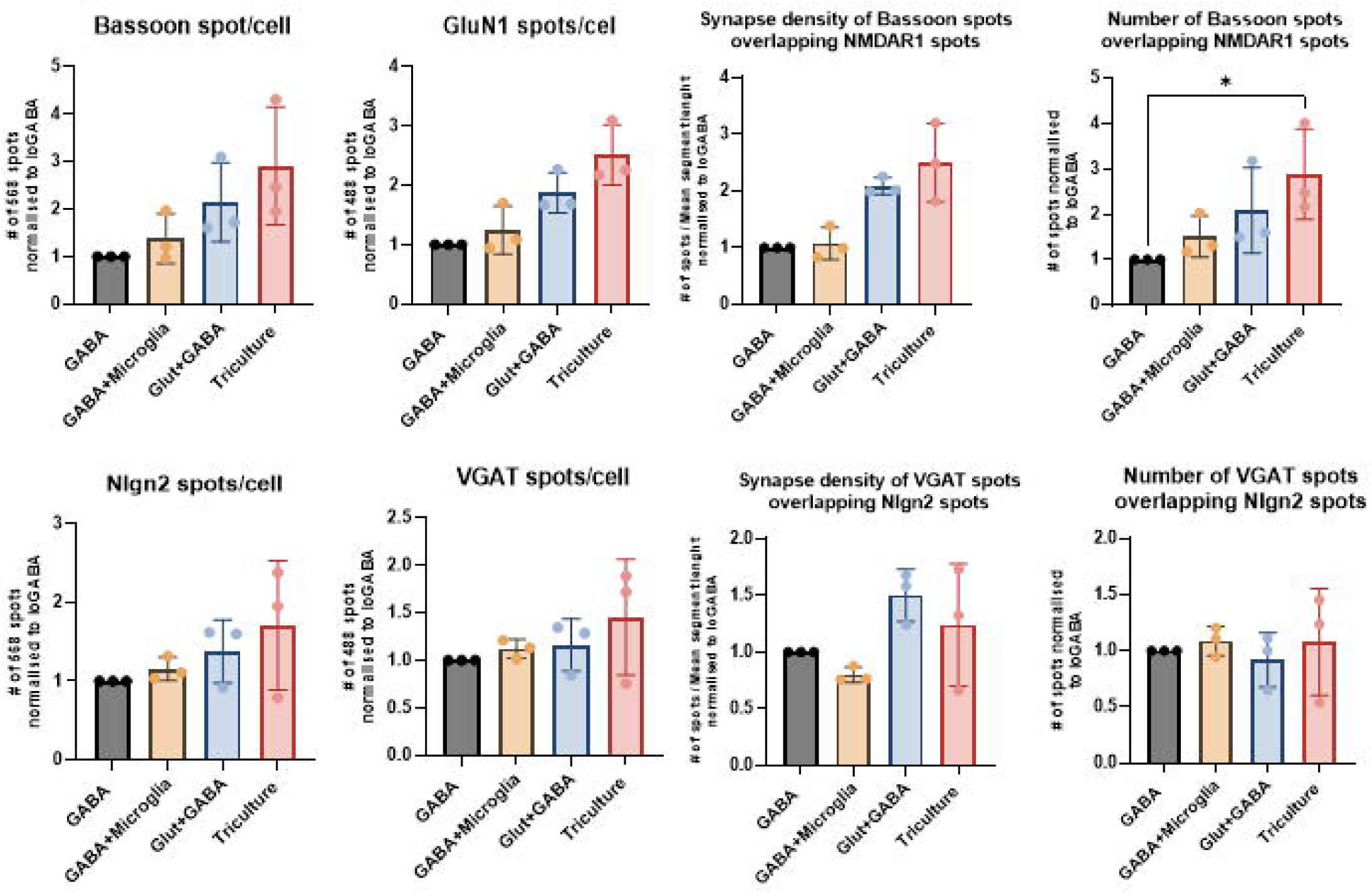
GABAergic neurons correctly form synapses in presence of microglia. **A)** Excitatory and **B)** Inhibitory synapses analysis on ioGABAergic (GABA) neurons in the presence and absence of microglia, normalised to GABA only condition (Kruskal-Wallis test, with Dunn’s post hoc test, N=3).

Overall, our data demonstrates that the tri-culture reliably forms both excitatory and inhibitory synapses and that microglia incorporation does not appear to alter early synaptogenesis under the conditions tested. Moreover, our automated analysis pipeline is a robust analytical framework for high-content quantification of synapses.

### Microglia incorporation increases network activity compared to neuronal cultures without microglia

To test the electrophysiological maturation of the tri-culture model, we seeded different culture conditions onto MEAs and monitored network properties longitudinally (Pavlinek *et al*., 2026) (**Figure 4A**). From spontaneous neuronal activity recordings, we extracted several complementary metrics describing neuronal firing and network behaviour (Pavlinek *et al*., 2026). Mean firing rate (MFR) and weighted mean firing rate (WMFR) quantify overall neuronal firing activity, with WMFR accounting only for electrodes that are electrically active. Burst number was used to assess structured neuronal firing patterns, while the synchrony index quantifies the temporal coordination of spikes across multiple electrodes, thus representing a network-level measure of functional connectivity and network maturation. We specifically measured these parameters in cultures comprised of ioGlutamatergic neurons alone, ioGlutamatergic+ioGABAergic (80:20 ratio), ioGlutamatergic+ioMicroglia, and ioGlutamatergic+ioGABAergic+ioMicroglia (tri-culture) conditions. Where present, microglia were added to DIV10 neurons.

**Figure 4.**
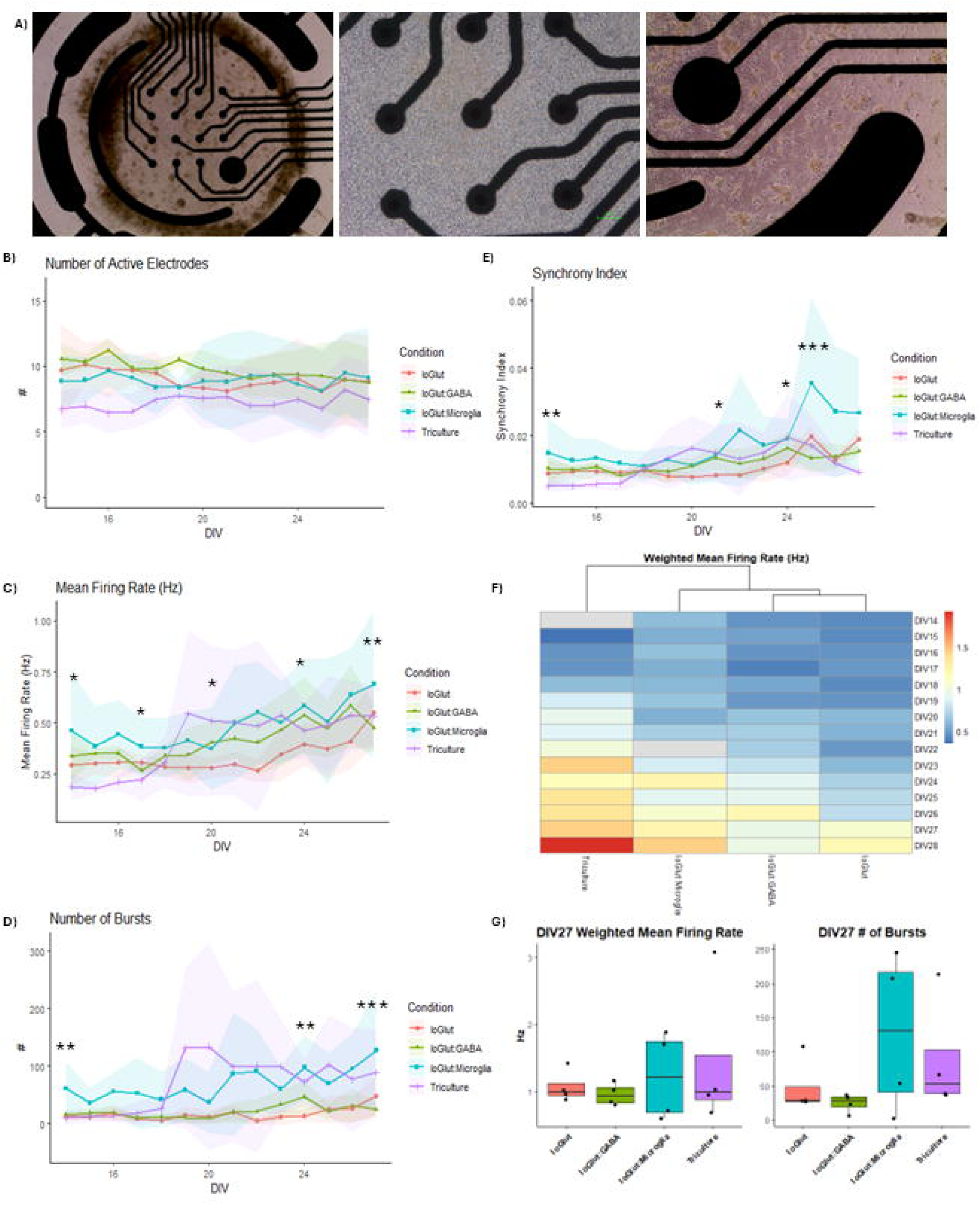
Neurons in the tri-culture model have more bursts compared to co-culture. **A)** Representative images of the triculture model on multi-well MEA plates, imaged with 4x, 10x and 20x brightfield objectives. **B)** Line graph of the Number of Active Electrodes, **C)** Mean Firing Rate, **D)** Number of Burst, and **E)** Synchrony Index over time, mean and standard deviation, Aligned Rank Transformation (ART) and Pairwise Comparisons, N=4. **F)** Heatmap showing the Weighted MFR over time, across conditions, **G)** Boxplots showing Weighted Mean Firing Rate and Number of Bursts at DIV27.

Network activities were recorded for 10 minutes daily across all cultures from DIV14 to DIV27. The number of active electrodes remained stable over time, indicating preserved network integrity and absence of cell loss (**Figure 4B**). As expected, MRF increased over time across all cultures, indicative of network maturation (**Figure 4C**). Comparisons across all culture conditions however, revealed several differences. In particular, ioGlutamatergic+ioMicroglia cultures exhibited increased MFR compared to ioGlutamatergic cultures at multiple timepoints. (**Figure 4C**). In contrast, tri-cultures displayed reduced MFR at early stages of network maturation, and comparable activity in later timepoints (**Figure 4C**).

Analysis of burst activity revealed that both ioGlut+ioMicroglia and tri-culture conditions displayed increased number of bursts beginning around DIV19 compared to ioGlut and ioGlut+ioGABA cultures (**Figure 4D**, **4G**). This indicates that the presence of microglia may promote the earlier emergence of structured firing patterns. In addition, ioGlut+ioMicroglia cultures exhibited a significantly higher synchrony index after DIV24 relative to control cultures, indicating increased coordination of activity across electrodes. In contrast, synchrony levels in the tri-culture remained comparable to neuronal mono- and co-cultures (**Figure 4E**), suggesting that the presence of inhibitory neurons may stabilize network-wide synchronization while still permitting increased burst activity.

Interestingly, WMFR, demonstrated the progressive maturation of network activity across all culture conditions (**Figure 4F–G**), with optimal firing for the tri-culture condition. Overall, the addition of microglia led to increased variability across several electrophysiological measures compared to neuronal cultures alone. Indeed, at DIV27, cultures containing microglia displayed greater variability in WMFR and burst number compared to cultures without microglia (**Figure 4G**).

Taken together, these results demonstrate that tri-cultures display stable electrophysiological maturation and can be used for longitudinal network analysis. While the presence of microglia increases variability in network activity, our data suggest that microglia modulate neuronal firing patterns during network maturation, while inclusion of inhibitory neurons helps maintain balanced network synchronisation.

### *TREM2* mutant microglia influence the burst phenotype but not the synapse density in early development

To assess whether the triculture platform could be used for disease modelling, we next tested its ability to detect functional effects of microglia carrying disease-associated genetic variants. As a proof-of-concept, we focused on *TREM2*, a microglial receptor known to regulate microglial activation and synapse-related processes for microglia activation and synaptic stability, and implicated in Alzheimer’s disease risk (Wang *et al*., 2015; Filipello *et al*., 2018; Li *et al*., 2023; Tagliatti *et al*., 2024; Feiten *et al*., 2026). To this end, we incorporated ioMicroglia carrying the Alzheimer’s disease-associated mutation *TREM2* R47H/R47H and compared them with wild-type (WT) microglia within the triculture system (Figure 5).

**Figure 5.**
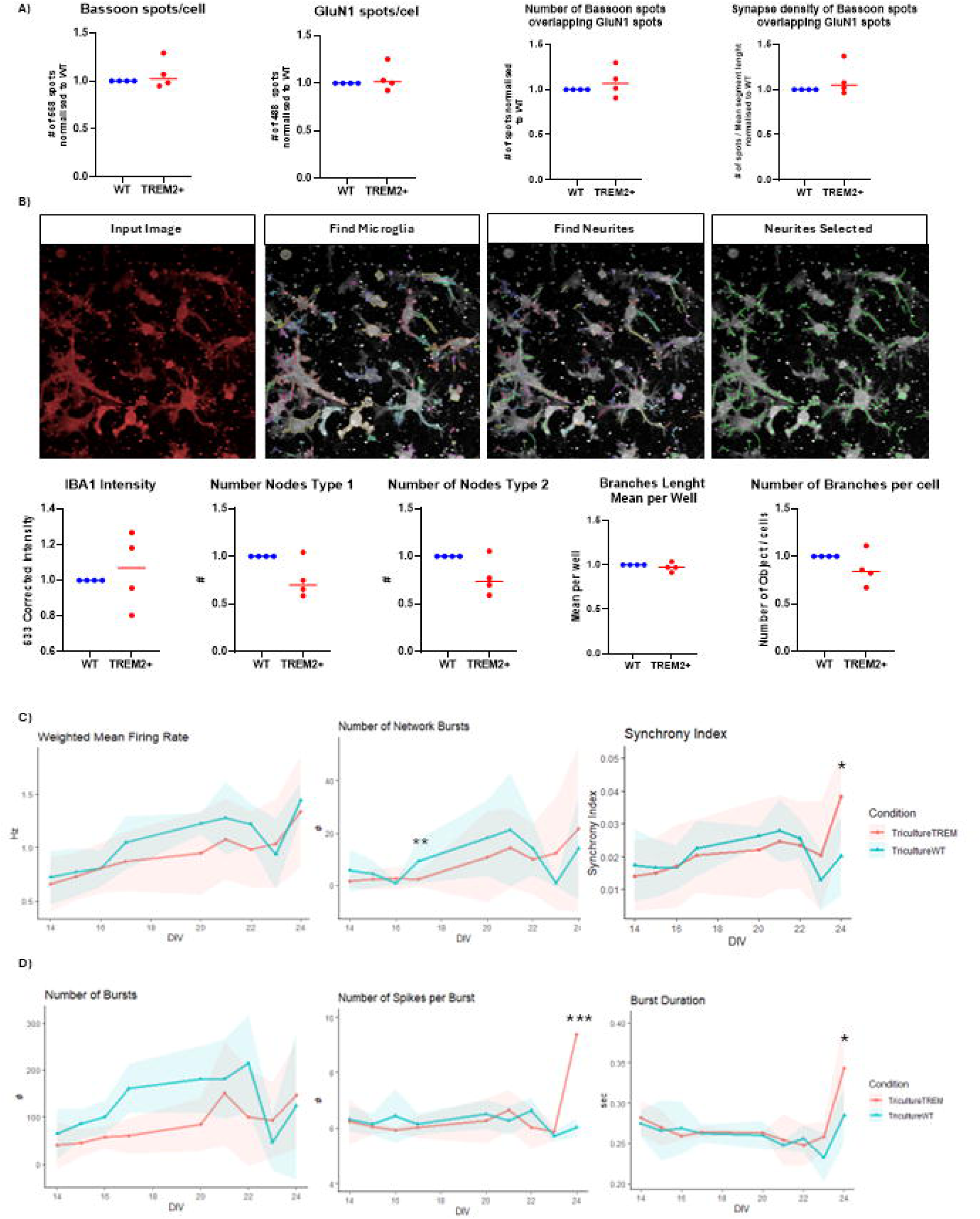
Triculture can be used to study microglia-specific genes and their phenotype. **A)** Excitatory synapses analysis in WT compared to mutant microglia. n=4, with 3 technical replicates. One-sample t-test. **B)** Overview of the imaging analysis pipeline developed to analyse microglia morphology and results of IBA1 intensity analysis and morphological analysis of mutant microglia compared to WT. N=4, with 3 technical replicates each. One-sample t-test. **C)** Line graph showing WMFR and Number of Network Bursts, average and standard deviation. **D)** Line graph showing number of Bursts, Spikes per bursts, and burst duration, average and standard deviation, Aligned Rank Transformation (ART), with Pairwise Comparisons.

We first assessed whether mutant microglia altered synaptogenesis of excitatory synapses in the tri-culture. Synapse analysis at DIV21 revealed no significant differences in either excitatory synaptic protein expression or synapse number between WT and *TREM2*-mutant microglia tri-cultures (**Figure 5A**). Next, we examined whether the *TREM2* R47H/R47H (io1035, bit.bio) affected microglial morphology within the tri-culture. To quantify morphological features, we developed an automated analysis pipeline that identifies microglia in the IBA1 channel and detects cellular branches using a neurite-detection algorithm (Figure 5B). Quantification of IBA1 expression (intensity), branch number, branch length, and node counts revealed no significant differences between WT and *TREM2*-mutant microglia (**Figure 5B**).

We further examined if *TREM2*-mutant microglia influenced neuronal network activity. MEA recordings were performed from DIV14 to DIV25 across tri-cultures containing WT or mutant *TREM2* containing microglia. Examination of network properties revealed that WMFR did not differ significantly between WT and mutant tricultures, suggesting that overall neuronal firing activity was unaffected (**Figure 5C**). However, at later timepoints, we observed abnormal bursting, characterised by a higher number of spikes and increased bust duration (**Figure 5C**). At DIV24, an increase in Synchrony Index is also seen in the conditions with mutant microglia. (**Figure 5D**).

Taken together, these data indicate that, under the conditions tested, while *TREM2* R47H mutant containing microglia do not alter synapse structure or microglia morphology, they subtly impact network burst properties. These results highlight the applicability of the tri-culture system to investigate and detect morphological and functional similarities and differences in a functional genomics context. They further demonstrate how this platform can be used to model microglia-driven mechanisms in neurological diseases.

## Discussion

Human induced pluripotent stem cell (hiPSC)-derived neuronal models are increasingly being used to study neuronal development, neural function, and disease mechanisms. Yet there remains a need for rapid and reproducible culture systems (Polit *et al*., 2023) with defined cellular composition to enable controlled investigation of neuron-glia interactions. Multiple lines of evidence suggest that human-derived models provide important platforms to study neuron-glia interactions and human-specific microglia biology (Abud *et al*., 2017; Popova *et al*., 2021; Cerneckis & Shi, 2023; Cserép *et al*., 2025; Woolf *et al*., 2025). Here, we present a scalable and consistent human tri-culture model comprising excitatory neurons, inhibitory neurons, and microglia. Our systematic characterisation of the tri-culture demonstrates that this system supports the structural and functional interrogation of microglia into co-cultures of excitatory and inhibitory neurons. Moreover, we showed that the presence of microglia in deterministically programmed neural networks impact the overall activity and synchronicity of neuronal cultures, consistent with recent findings (Mordelt *et al*., 2026). We further show that this tri-culture system can be used to assess how microglia harbouring disease-relevant mutations influence synapse structure and neuronal network function. Taken together, by combining deterministically-programmed cell types with controlled cellular ratios, this system provides a tractable platform for investigating how microglia influence neuronal network development under physiological and disease-relevant conditions.

A key feature of this tri-culture model is the inclusion of both glutamatergic and GABAergic neurons in a defined 80:20 ratio prior to microglial integration. This design was motivated by growing evidence that E/I balance can shape synapse structure, network function and pharmacological responsiveness (Mossink *et al*., 2021). Alterations in E/I balance is thought to be a major contributor to the pathophysiology of psychiatric and neurodegenerative disorders (Mossink *et al*., 2021; Chen *et al*., 2025). While several recent human *in vitro* neuroimmune models have incorporated glial cell types to increase biological complexity (Pascual *et al*., 2012; Lish *et al*., 2025a), many rely on excitatory neuronal mono-cultures or do not explicitly control inhibitory composition. Previous work using hiPSC-derived neuronal networks has demonstrated that heterogeneous E/I cultures exhibit more physiologically relevant and structured bursting activity compared to homogeneous excitatory or inhibitory networks, highlighting the importance of defined E/I balance for network maturation and stability (Mossink *et al*., 2021; Gu *et al*., 2026). Our results show that optimisation of neuronal ratios is important for generating a stable and reproducible network function. Consistently with these observations, 80:20 E/I compositions have been shown to support coordinated and synchronized network activity, while both purely excitatory and highly inhibitory configurations display less organised dynamics (Crocco *et al*., 2025). In addition, inhibitory neurons have been shown to regulate burst timing and structure, while excitatory neurons contribute to burst amplitude, indicating that E/I balance governs complementary aspects of network activity (Parodi *et al*., 2023). Thus, in this context, the incorporation of microglia into a culture with defined E/I ratio that approximates physiological cortical ratios *in vivo* (Cerina *et al*., 2023), provides a controlled framework in which to examine microglial effects on neuronal activity.

The characterisation of synapses demonstrated that the co- and tri-cultures reliably form both excitatory and inhibitory synaptic structures. Interestingly, incorporation of microglia does not disrupt early synaptogenesis under the conditions tested. Although microglia are well established regulators of synapse remodelling and pruning (Ji *et al*., 2013; Sellgren *et al*., 2019; Andoh & Koyama, 2021; Faust *et al*., 2025), we did not detect significant changes in synaptic density at DIV21. This likely reflects the developmental window examined and is in line with previous reports (Hume, 2025; O’Keeffe *et al*., 2025). Indeed, major pruning-related effects may emerge later or require additional maturation and environmental cues (Paolicelli *et al*., 2011; Couch *et al*., 2024). Despite the absence of an effect on synapse number, analysis of network function revealed an increase in overall activity, consistent with existing data (Kok *et al*., 2025), meaning our tri-culture model can replicate reproducible phenotype.

Finally, we proved an example on how the model can be applied to investigate disease-relevant genetic variants. Here we challenged the tri-culture model with *TREM2* R47H/R47H microglia. *TREM2* is a key microglial receptor implicated in metabolic fitness, activation state, and Alzheimer’s disease risk (Ulland & Colonna, 2018) (Wang *et al*., 2015) (Li *et al*., 2023). It has also been linked to synapse-related phenotypes in experimental systems (Filipello *et al*., 2018). However, published evidence of *TREM2* involved in synaptic regulation have mostly been obtained from murine models (Filipello *et al*., 2018; Penney *et al*., 2023). In our human tri-culture model, the presence of *TREM2* R47H mutation carrying microglia did not alter excitatory synapse markers or gross microglial morphology, but it did produce subtle yet reproducible changes in burst-related network properties, including early increases in burst and network burst frequency and later changes in burst structure. Of note, similar results were previously reported using murine *TREM2*-/-hippocampal neurons (Tagliatti *et al*., 2024), and within cortical brain regions of TREM2 mutant mice (Das *et al*., 2023; Johnston *et al*., 2024) including alterations in neuronal networks. These findings are important for two reasons. First, they suggest that microglial genetic perturbation can influence neuronal network behaviour even in the absence of overt early structural phenotypes. Second, they demonstrate that the tri-culture system is sufficiently sensitive to detect disease-relevant functional effects of microglial variants in a human-defined neuronal environment.

It is important to note the limitations of this study. The model captures an early developmental stage and may therefore underestimate later-arising effects on synapse remodelling or microglial maturation. In addition, the system is two-dimensional and does not include astrocytes, which are likely to influence both synaptic development and inflammatory signalling. However, this reduced complexity is also a strength, as it allows focused analysis of microglia-specific effects within a controlled and reproducible neuronal network. Future work could extend the maturation window, incorporate additional glial components, or apply the platform to patient-derived and engineered microglial variants relevant to neurodevelopmental and neurodegenerative disorders.

Overall, our work demonstrates a rapid, defined, reproducible and scalable human neuron-microglia tri-culture model that integrates physiological E/I balance with structural and functional network readouts. The platform supports robust structural and electrophysiological characterisation and is sensitive to disease-relevant microglial perturbation. As such, it provides a useful intermediate system between simple neuronal mono-cultures and more complex multicellular models for studying how microglia shape human neuronal network development and dysfunction.

## ACKNOWLEDGMENTS

The authors acknowledge funding support from UK Medical Research Council, Grant Nos. MR/Y012968/1, MR/Y008170/1 and MR/Y012968/1 (A.C.V., D.P.S.). This work was supported by bit.bio through contribution to the research funding and in□Ikind provision of ioCells products. D.P.S. is also a recipient of an Independent Researcher Award from the Brain and Behavior Foundation (Grant No. 25957). A.P. is in receipt of the MRC- Institute for Translational Neurodevelopment (ITND) Ph.D. studentship, as part of the MRC Centre for Neurodevelopmental Disorders (Medical Research Council MR/P502108/1). The authors thank the Wohl Cellular Imaging Centre (King’s College London) for technical support. Schematic diagrams in Figures 1 and 2 were created using BioRender.

## AUTHOR CONTRIBUTIONS

Conceptualization, A.C.V. and D.P.S.; methodology, S.G., A.P., O.L.C., A.C.V. and D.P.S.; Investigation, S.G., O.L.C., G.C.; writing-original draft, S.G., A.C.V. and D.P.S.; visualization, S.G., writing-review & editing, S.G., A.C.V. and D.P.S.; funding acquisition, A.C.V. and D.P.S.; supervision, A.C.V., and D.P.S.

## References

Abud, E.M., Ramirez, R.N., Martinez, E.S., Healy, L.M., Nguyen, C.H.H., Newman, S.A., Yeromin, A.V., Scarfone, V.M., Marsh, S.E., Fimbres, C., Caraway, C.A., Fote, G.M., Madany, A.M., Agrawal, A., Kayed, R., Gylys, K.H., Cahalan, M.D., Cummings, B.J., Antel, J.P., Mortazavi, A., Carson, M.J., Poon, W.W. & Blurton-Jones, M. (2017) iPSC-Derived Human Microglia-like Cells to Study Neurological Diseases. Neuron, 94, 278–293 e279.

Andoh, M. & Koyama, R. (2021) Microglia regulate synaptic development and plasticity. Developmental Neurobiology, 81.

Badimon, A., Strasburger, H.J., Ayata, P., Chen, X., Nair, A., Ikegami, A., Hwang, P., Chan, A.T., Graves, S.M., Uweru, J.O., Ledderose, C., Kutlu, M.G., Wheeler, M.A., Kahan, A., Ishikawa, M., Wang, Y.-C., Loh, Y.-H.E., Jiang, J.X., Surmeier, D.J., Robson, S.C., Junger, W.G., Sebra, R., Calipari, E.S., Kenny, P.J., Eyo, U.B., Colonna, M., Quintana, F.J., Wake, H., Gradinaru, V., Schaefer, A., Badimon, A., Strasburger, H.J., Ayata, P., Chen, X., Nair, A., Ikegami, A., Hwang, P., Chan, A.T., Graves, S.M., Uweru, J.O., Ledderose, C., Kutlu, M.G., Wheeler, M.A., Kahan, A., Ishikawa, M., Wang, Y.-C., Loh, Y.-H.E., Jiang, J.X., Surmeier, D.J., Robson, S.C., Junger, W.G., Sebra, R., Calipari, E.S., Kenny, P.J., Eyo, U.B., Colonna, M., Quintana, F.J., Wake, H., Gradinaru, V. & Schaefer, A. (2020) Negative feedback control of neuronal activity by microglia. Nature 2020 586:7829, 586.

Beninger, J., Rossbroich, J., Tóth, K. & Naud, R. (2024) Functional subtypes of synaptic dynamics in mouse and human. Cell Reports, 43.

Berryer, M.H., Rizki, G., Nathanson, A., Klein, J.A., Trendafilova, D., Susco, S.G., Lam, D., Messana, A., Holton, K.M., Karhohs, K.W., Cimini, B.A., Pfaff, K., Carpenter, A.E., Rubin, L.L. & Barrett, L.E. (2023) High-content synaptic phenotyping in human cellular models reveals a role for BET proteins in synapse assembly. eLife, 12.

Breschi, A., Gingeras, T.R. & Guigó, R. (2017) Comparative transcriptomics in human and mouse. Nature reviews. Genetics, 18.

Cerina, M., Piastra, M.C., Frega, M., Cerina, M., Piastra, M.C. & Frega, M. (2023) The potential of in vitro neuronal networks cultured on micro electrode arrays for biomedical research. Progress in Biomedical Engineering, 5.

Cerneckis, J. & Shi, Y. (2023) Context matters: hPSC-derived microglia thrive in a humanized brain environment in vivo. Cell stem cell, 30.

Chen, Z.-P., Zhao, X., Wang, S., Cai, R., Liu, Q., Ye, H., Wang, M.-J., Peng, S.-Y., Xue, W.-X., Zhang, Y.-X., Li, W., Tang, H., Huang, T., Zhang, Q., Li, L., Gao, L., Zhou, H., Hang, C., Zhu, J.-N., Li, X., Liu, X., Cong, Q., Yan, C., Chen, Z.-P., Zhao, X., Wang, S., Cai, R., Liu, Q., Ye, H., Wang, M.-J., Peng, S.-Y., Xue, W.-X., Zhang, Y.-X., Li, W., Tang, H., Huang, T., Zhang, Q., Li, L., Gao, L., Zhou, H., Hang, C., Zhu, J.-N., Li, X., Liu, X., Cong, Q. & Yan, C. (2025) GABA-dependent microglial elimination of inhibitory synapses underlies neuronal hyperexcitability in epilepsy. Nature Neuroscience 2025 28:7, 28.

Couch, A.C.M., Brown, A.M., Raimundo, C., Solomon, S., Taylor, M., Sichlinger, L., Matuleviciute, R., Srivastava, D.P. & Vernon, A.C. (2024) Transcriptional and cellular response of hiPSC-derived microglia-neural progenitor co-cultures exposed to IL-6. Brain, Behavior, and Immunity, 122.

Crocco, E., Iannello, L., Tonelli, F., Lagani, G., Pandolfini, L., Ferro, M., Amato, G., Garbo, A.D. & Cremisi, F. (2025) A proper excitatory/inhibitory ratio is required to develop synchronized network activity in mouse cortical cultures. Stem Cell Reports, 20.

Cserép, C., Berki, P., Mars, M., Pósfai, B., Pasterkamp, R.J., Benkő, S. & Dénes, Á. (2025) Lessons from ex vivo and in vitro models in microglia research. Trends in Neurosciences, 48.

Das, M., Mao, W., Voskobiynyk, Y., Necula, D., Lew, I., Petersen, C., Zahn, A., Yu, G.-Q., Yu, X., Smith, N., Sayed, F.A., Gan, L., Paz, J.T. & Mucke, L. (2023) Alzheimer risk-increasing TREM2 variant causes aberrant cortical synapse density and promotes network hyperexcitability in mouse models. Neurobiology of Disease, 186.

Deng, B. (2017) Mouse models and induced pluripotent stem cells in researching psychiatric disorders. Stem Cell Investigation, 4.

Eyo, U. & Molofsky, A.V. (2023) Defining microglial-synapse interactions. Science, 381.

Faust, T.E., Lee, Y.-H., O’Connor, C.D., Boyle, M.A., Gunner, G., Durán-Laforet, V., Ferrari, L.L., Murphy, R.E., Badimon, A., Sakers, K., Eroglu, C., Ayata, P., Schaefer, A. & Schafer, D.P. (2025) Microglia-astrocyte crosstalk regulates synapse remodeling via Wnt signaling. Cell, 188.

Feiten, A.F., Dahm, K., Schlepckow, K., van Lengerich, B., Suh, J.H., Reifschneider, A., Wefers, B., Bartos, L.M., Wind-Mark, K., de Weerd, L., Ulas, T., De-Domenico, E., Grundschöttel, P., Paulusch, S., Tast, B., Honda, T., Müller, S.A., Becker, M., Khalin, I., Ricci, A., Liesz, A., Brunner, B., Krenner, C., Buschmann, K., Nuscher, B., Spieth, L., Junker, N., Berghoff, S.A., Davis, S.S., Neher, J.J., Wurst, W., Plesnila, N., Lewcock, J.W., Simons, M., Lichtenthaler, S.F., Di Paolo, G., Brendel, M., Capell, A., Monroe, K.M., Schultze, J.L., Haass, C., Feiten, A.F., Dahm, K., Schlepckow, K., van Lengerich, B., Suh, J.H., Reifschneider, A., Wefers, B., Bartos, L.M., Wind-Mark, K., de Weerd, L., Ulas, T., De-Domenico, E., Grundschöttel, P., Paulusch, S., Tast, B., Honda, T., Müller, S.A., Becker, M., Khalin, I., Ricci, A., Liesz, A., Brunner, B., Krenner, C., Buschmann, K., Nuscher, B., Spieth, L., Junker, N., Berghoff, S.A., Davis, S.S., Neher, J.J., Wurst, W., Plesnila, N., Lewcock, J.W., Simons, M., Lichtenthaler, S.F., Di Paolo, G., Brendel, M., Capell, A., Monroe, K.M., Schultze, J.L. & Haass, C. (2026) TREM2 expression level is critical for microglial state, metabolic capacity and efficacy of TREM2 agonism. Nature Communications 2026 17:1, 17.

Filipello, F., Morini, R., Corradini, I., Zerbi, V., Canzi, A., Michalski, B., Erreni, M., Markicevic, M., Starvaggi-Cucuzza, C., Otero, K., Piccio, L., Cignarella, F., Perrucci, F., Tamborini, M., Genua, M., Rajendran, L., Menna, E., Vetrano, S., Fahnestock, M., Paolicelli, R.C. & Matteoli, M. (2018) The Microglial Innate Immune Receptor TREM2 Is Required for Synapse Elimination and Normal Brain Connectivity. Immunity, 48.

Gu, J., Rollo, B., Liu, Z., O’Brien, T.J., Kwan, P., Cromer, B. & Sumer, H. (2026) Forward programming of human pluripotent stem cells to generate glutamatergic and GABAergic neurons in a tri-culture model with astrocytes. Stem Cell Res Ther.

Habich, C., Miller, L.N., Luz, N.G., Malik, L.I., Chegireddy, K., Maneshi, M.M., Wolf, M., Kummer, M., Iriki, H., Röwe, J., Marro, S., Scott, V.E., Smith, K.M., Kim, B.S., Reinhardt, P. & Goedken, E.R. (2025) NGN2 Expression and Regional Patterning Allow Rapid Differentiation from hiPSCs to DRG-Like Neurons Responsive to Type 2 Cytokines. bioRxiv.

Hanger, B., Couch, A., Rajendran, L., Srivastava, D.P. & Vernon, A.C. (2020) Emerging Developments in Human Induced Pluripotent Stem Cell-Derived Microglia: Implications for Modelling Psychiatric Disorders With a Neurodevelopmental Origin. Front Psychiatry, 11, 789.

Haskell, A.K., Kulas, J.A., Carter, W.E., Javens-Wolfe, J., Hinkel, R.D., Moussaif, M., Smiley, J.S., Lazaro, O., Robertson, S., Palkowitz, A.D., Lamb, B.T., Richardson, T.I., Dage, J.L., Chu, S., Johnson, T., Stancato, L.F. & Syed, A.Q. (2026) Generation and characterization of iPSC-derived microglia for in vitro modeling of stimuli-specific neuroimmune responses. Alzheimer’s & Dementia, 22.

Hasselmann, J. & Blurton-Jones, M. (2020) Human iPSC-derived microglia: A growing toolset to study the brain’s innate immune cells. Glia, 68.

Hume, D.A. (2025) Life without microglia. Trends in Neurosciences, 48.

Ji, K., Akgul, G., Wollmuth, L.P. & Tsirka, S.E. (2013) Microglia Actively Regulate the Number of Functional Synapses. PLOS ONE, 8.

Johnston, K.G., Berackey, B.T., Tran, K.M., Gelber, A., Yu, Z., MacGregor, G.R., Mukamel, E.A., Tan, Z., Green, K.N., Xu, X., Johnston, K.G., Berackey, B.T., Tran, K.M., Gelber, A., Yu, Z., MacGregor, G.R., Mukamel, E.A., Tan, Z., Green, K.N. & Xu, X. (2024) Single-cell spatial transcriptomics reveals distinct patterns of dysregulation in non-neuronal and neuronal cells induced by the Trem2R47H Alzheimer’s risk gene mutation. Molecular Psychiatry 2024 30:2, 30.

Kok, L.M.L., Helwegen, K., Coveña, N.F. & Heine, V.M. (2025) Human pluripotent stem cell-derived microglia shape neuronal morphology and enhance network activity in vitro. Journal of Neuroscience Methods, 415.

Kuhn, S.A., Landeghem, F.K.H.v., Zacharias, R., Färber, K., Rappert, A., Pavlovic, S., Hoffmann, A., Nolte, C. & Kettenmann, H. (2004) Microglia express GABAB receptors to modulate interleukin release. Molecular and Cellular Neuroscience, 25.

Li, E., Benitez, C., Boggess, S.C., Koontz, M., Rose, I.V.L., Martinez, D., Dräger, N., Teter, O.M., Samelson, A.J., Pierce, N.i., Ullian, E.M. & Kampmann, M. (2025) CRISPRi-based screens in iAssembloids to elucidate neuron-glia interactions. Neuron, 113.

Li, Y., Xu, H., Wang, H., Yang, K., Luan, J. & Wang, S. (2023) TREM2: Potential therapeutic targeting of microglia for Alzheimer’s disease. Biomedicine & Pharmacotherapy, 165.

Lish, A.M., Ashour, N., Pearse, R.V., 2nd, Galle, P.C., Orme, G.A., Heuer, S.E., Benoit, C.R., Alexander, K.D., Grogan, E.F.L., Terzioglu, G., Scarpa, A., Stern, A.M., Seyfried, N., Menon, V. & Young-Pearse, T.L. (2025a) Astrocyte induction of disease-associated microglia is suppressed by acute exposure to fAD neurons in human iPSC triple cultures. Cell Rep, 44, 115777.

Lish, A.M., Galle, P.C., Orme, G.A., Ashour, N., Heuer, S.E., Curle, A.J., Muratore, C.R. & Young-Pearse, T.L. (2025b) Protocol for generating a human iPSC-derived tri-culture model to study interactions between neurons, astrocytes, and microglia. STAR Protocols, 6.

Luchena, C., Zuazo-Ibarra, J., Valero, J., Matute, C., Alberdi, E. & Capetillo-Zarate, E. (2022) Frontiers | A Neuron, Microglia, and Astrocyte Triple Co-culture Model to Study Alzheimer’s Disease. Frontiers in Aging Neuroscience, 14.

Marin, O. (2025) Development of GABAergic Interneurons in the Human Cerebral Cortex. Eur J Neurosci, 61, e70136.

Matuleviciute, R., Akinluyi, E.T., Muntslag, T.A.O., Dewing, J.M., Long, K.R., Vernon, A.C., Tremblay, M.-E. & Menassa, D.A. (2023) Microglial contribution to the pathology of neurodevelopmental disorders in humans. Acta Neuropathologica, 146, 663–683.

Mondelli, V., Vernon, A.C., Turkheimer, F., Dazzan, P. & Pariante, C.M. (2017) Brain microglia in psychiatric disorders. Lancet Psychiatry, 4, 563–572.

Mordelt, A., Scheefhals, N., Schuurmans, I.M.E., Slottje, K., Rivera, M.C., Hommersom, M.P., Huang, A., Bichmann, L., Lewerissa, E.I., Hugte, E.J.H.v., Schubert, D., Tsang, J.S., Kasri, N.N. & Witte, L.D.d. (2026) MEA-LINK identifies the CCL4-CCR5 axis in neuronal hyperactivity control by human microglia. bioRxiv.

Mordelt, A., Schuurmans, I.M.E., Scheefhals, N., Hommersom, M.P., Slottje, K., Mast, K., Graziani, M., González, C.O., Wingens, L.J.A., Schubert, D., Kasri, N.N. & Witte, L.D.d. (2025) Long-term co-maturation of stem cell-derived microglia and neuronal networks: an optimized platform to assess human microglial contribution to neuronal function. bioRxiv.

Mossink, B., van Rhijn, J.-R., Wang, S., Linda, K., Vitale, M.R., Zöller, J.E.M., van Hugte, E.J.H., Bak, J., Verboven, A.H.A., Selten, M., Negwer, M., Latour, B.L., van der Werf, I., Keller, J.M., Klein Gunnewiek, T.M., Schoenmaker, C., Oudakker, A., Anania, A., Jansen, S., Lesch, K.-P., Frega, M., van Bokhoven, H., Schubert, D., Nadif Kasri, N., Mossink, B., van Rhijn, J.-R., Wang, S., Linda, K., Vitale, M.R., Zöller, J.E.M., van Hugte, E.J.H., Bak, J., Verboven, A.H.A., Selten, M., Negwer, M., Latour, B.L., van der Werf, I., Keller, J.M., Klein Gunnewiek, T.M., Schoenmaker, C., Oudakker, A., Anania, A., Jansen, S., Lesch, K.-P., Frega, M., van Bokhoven, H., Schubert, D. & Nadif Kasri, N. (2021) Cadherin-13 is a critical regulator of GABAergic modulation in human stem-cell-derived neuronal networks. Molecular Psychiatry 2021 27:1, 27.

Mossink, B., van Rhijn, J.R., Wang, S., Linda, K., Vitale, M.R., Zoller, J.E.M., van Hugte, E.J.H., Bak, J., Verboven, A.H.A., Selten, M., Negwer, M., Latour, B.L., van der Werf, I., Keller, J.M., Klein Gunnewiek, T.M., Schoenmaker, C., Oudakker, A., Anania, A., Jansen, S., Lesch, K.P., Frega, M., van Bokhoven, H., Schubert, D. & Nadif Kasri, N. (2022) Cadherin-13 is a critical regulator of GABAergic modulation in human stem-cell-derived neuronal networks. Mol Psychiatry, 27, 1–18.

Nayak, D., Roth, T.L. & McGavern, D.B. (2014) Microglia Development and function. Annual review of immunology, 32.

Nieland, T.J.F., Logan, D.J., Saulnier, J., Lam, D., Johnson, C., Root, D.E., Carpenter, A.E. & Sabatini, B.L. (2014) High Content Image Analysis Identifies Novel Regulators of Synaptogenesis in a High-Throughput RNAi Screen of Primary Neurons. PLOS ONE, 9.

O’Keeffe, M., Booker, S.A., Walsh, D., Li, M., Henley, C., Simões de Oliveira, L., Liu, M., Wang, X., Banqueri, M., Ridley, K., Dissanayake, K.N., Martinez-Gonzalez, C., Craigie, K.J., Vasoya, D., Leah, T., He, X., Hume, D.A., Duguid, I., Nolan, M.F., Qiu, J., Wyllie, D.J.A., Dando, O.R., Gonzalez-Sulser, A., Gan, J., Pridans, C., Kind, P.C., Hardingham, G.E., O’Keeffe, M., Booker, S.A., Walsh, D., Li, M., Henley, C., Simões de Oliveira, L., Liu, M., Wang, X., Banqueri, M., Ridley, K., Dissanayake, K.N., Martinez-Gonzalez, C., Craigie, K.J., Vasoya, D., Leah, T., He, X., Hume, D.A., Duguid, I., Nolan, M.F., Qiu, J., Wyllie, D.J.A., Dando, O.R., Gonzalez-Sulser, A., Gan, J., Pridans, C., Kind, P.C. & Hardingham, G.E. (2025) Typical development of synaptic and neuronal properties can proceed without microglia in the cortex and thalamus. Nature Neuroscience 2025 28:2, 28.

Odawara, A., Saitoh, Y., Alhebshi, A.H., Gotoh, M. & Suzuki, I. (2014) Long-term electrophysiological activity and pharmacological response of a human induced pluripotent stem cell-derived neuron and astrocyte co-culture. Biochem Biophys Res Commun, 443, 1176–1181.

Paolicelli, R.C., Bolasco, G., Pagani, F., Maggi, L., Scianni, M., Panzanelli, P., Giustetto, M., Ferreira, T.A., Guiducci, E., Dumas, L., Ragozzino, D. & Gross, C.T. (2011) Synaptic Pruning by Microglia Is Necessary for Normal Brain Development. Science, 333.

Papandreou, A., Luft, C., Barral, S., Kriston-Vizi, J., Kurian, M.A. & Ketteler, R. (2023) Automated high-content imaging in iPSC-derived neuronal progenitors. SLAS Discovery, 28.

Parodi, G., Brofiga, M., Pastore, V.P., Chiappalone, M. & Martinoia, S. (2023) Deepening the role of excitation/inhibition balance in human iPSCs-derived neuronal networks coupled to MEAs during long-term development. J Neural Eng, 20.

Pascual, O., Achour, S.B., Rostaing, P., Triller, A., Bessis, A., Pascual, O., Ben Achour, S., Rostaing, P., Triller, A. & Bessis, A. (2012) Microglia activation triggers astrocyte-mediated modulation of excitatory neurotransmission. Proceedings of the National Academy of Sciences, 109.

Pavlinek, A., Guerrisi, S., O’Driscoll, K., Polit, L.D., Nagy, R., Lancaster, M.A., Vernon, A.C. & Srivastava, D.P. (2026) Electrophysiological development and functional plasticity in dissociated human cerebral organoids across multiple cell lines. Cell Reports Methods, 0.

Pawlowski, M., Ortmann, D., Bertero, A., Tavares, J.M., Pedersen, R.A., Vallier, L. & Kotter, M.R.N. (2017) Inducible and Deterministic Forward Programming of Human Pluripotent Stem Cells into Neurons, Skeletal Myocytes, and Oligodendrocytes. Stem Cell Reports, 8, 803–812.

Penney, J., Ralvenius, W.T., Loon, A., Cerit, O., Dileep, V., Milo, B., Pao, P.-C., Woolf, H. & Tsai, L.-H. (2023) iPSC-derived microglia carrying the TREM2 R47H/+ mutation are pro-inflammatory and promote synapse loss. Glia, 72.

Polit, L.D., Eidhof, I., McNeill, R.V., Warre-Cornish, K.M., Ohki, C.M.Y., Walter, N.M., Sala, C., Verpelli, C., Radtke, F., Galderisi, S., Mucci, A., Collo, G., Edenhofer, F., Castrén, M.L., Réthelyi, J.M., Ejlersen, M., Hohmann, S.S., Ilieva, M.S., Lukjanska, R., Matuleviciute, R. & Srivastava, D.P. (2023) Recommendations, guidelines, and best practice for the use of human induced pluripotent stem cells for neuropharmacological studies of neuropsychiatric disorders. Neuroscience Applied, 2.

Popova, G., Soliman, S.S., Kim, C.N., Keefe, M.G., Hennick, K.M., Jain, S., Li, T., Tejera, D., Shin, D., Chhun, B.B., McGinnis, C.S., Speir, M., Gartner, Z.J., Mehta, S.B., Haeussler, M., Hengen, K.B., Ransohoff, R.R., Piao, X. & Nowakowski, T.J. (2021) Human microglia states are conserved across experimental models and regulate neural stem cell responses in chimeric organoids. Cell Stem Cell, 28.

Que, Z., Olivero-Acosta, M.I., Robinson, M., Chen, I., Zhang, J., Wettschurack, K., Wu, J., Xiao, T., Otterbacher, C.M., Shankar, V., Harlow, H., Hong, S., Zirkle, B., Wang, M., Cui, N., Mandal, P., Chen, X., Deming, B., Halurkar, M., Zhao, Y., Rochet, J.-C., Xu, R., Brewster, A.L., Wu, L.-j., Yuan, C., Skarnes, W.C. & Yang, Y. (2024) Human IPSC-Derived Microglia Sense and Dampen Hyperexcitability of Cortical Neurons Carrying the Epilepsy-Associated SCN2A-L1342P Mutation. The Journal of Neuroscience, 45.

Rittenhouse, A., Krall, C., Plotkin, J., Alam El Din, D.M., Kincaid, B., Laird, J. & Smirnova, L. (2025) Microglia-containing neural organoids as brain microphysiological systems for long-term culture. Front Cell Neurosci, 19, 1616470.

Schafer, Dorothy P., Lehrman, Emily K., Kautzman, Amanda G., Koyama, R., Mardinly, Alan R., Yamasaki, R., Ransohoff, Richard M., Greenberg, Michael E., Barres, Ben A. & Stevens, B. (2012) Microglia Sculpt Postnatal Neural Circuits in an Activity and Complement-Dependent Manner. Neuron, 74.

Sellgren, C.M., Gracias, J., Watmuff, B., Biag, J.D., Thanos, J.M., Whittredge, P.B., Fu, T., Worringer, K., Brown, H.E., Wang, J., Kaykas, A., Karmacharya, R., Goold, C.P., Sheridan, S.D. & Perlis, R.H. (2019) Increased synapse elimination by microglia in schizophrenia patient-derived models of synaptic pruning. Nat Neurosci, 22, 374–385.

Sichlinger, L., Wulf, M., Polit, L.D., Nasser, F., Duarte, R.R.R., Powell, T.R., Marcus, K., Vernon, A.C. & Srivastava, D.P. (2024) Schizophrenia risk gene ZNF804A controls ribosome localization and synaptogenesis in developing human neurons. bioRxiv.

Summers, R.A., Fagiani, F., Rowitch, D.H., Absinta, M. & Reich, D.S. (2024) Novel human iPSC models of neuroinflammation in neurodegenerative disease and regenerative medicine. Trends in Immunology, 45.

Tagliatti, E., Desiato, G., Mancinelli, S., Bizzotto, M., Gagliani, M.C., Faggiani, E., Hernández-Soto, R., Cugurra, A., Poliseno, P., Miotto, M., Argüello, R.J., Filipello, F., Cortese, K., Morini, R., Lodato, S. & Matteoli, M. (2024) Trem2 expression in microglia is required to maintain normal neuronal bioenergetics during development. Immunity, 57.

Tegtmeyer, M., Liyanage, D., Han, Y., Hebert, K.B., Pei, R., Way, G.P., Ryder, P.V., Hawes, D., Tromans-Coia, C., Cimini, B.A., Carpenter, A.E., Singh, S., Nehme, R., Tegtmeyer, M., Liyanage, D., Han, Y., Hebert, K.B., Pei, R., Way, G.P., Ryder, P.V., Hawes, D., Tromans-Coia, C., Cimini, B.A., Carpenter, A.E., Singh, S. & Nehme, R. (2025) Combining phenomics with transcriptomics reveals cell-type-specific morphological and molecular signatures of the 22q11.2 deletion. Nature Communications 2025 16:1, 16.

Teter, O.M., McQuade, A., Hagan, V., Liang, W., Drager, N.M., Sattler, S.M., Holmes, B.B., Castillo, V.C., Papakis, V., Leng, K., Boggess, S., Nowakowski, T.J., Wells, J. & Kampmann, M. (2025) CRISPRi-based screen of autism spectrum disorder risk genes in microglia uncovers roles of ADNP in microglia endocytosis and synaptic pruning. Mol Psychiatry, 30, 4176–4193.

Ulland, T.K. & Colonna, M. (2018) TREM2 - a key player in microglial biology and Alzheimer disease. Nat Rev Neurol, 14, 667–675.

Upthegrove, R., Corsi-Zuelli, F., Couch, A.C.M., Barnes, N.M. & Vernon, A.C. (2025) Current Position and Future Direction of Inflammation in Neuropsychiatric Disorders: A Review. JAMA Psychiatry, 82, 1030–1046.

Wang, Y., Cella, M., Mallinson, K., Ulrich, J.D., Young, K.L., Robinette, M.L., Gilfillan, S., Krishnan, G.M., Sudhakar, S., Zinselmeyer, B.H., Holtzman, D.M., Cirrito, J.R. & Colonna, M. (2015) TREM2 lipid sensing sustains the microglial response in an Alzheimer’s disease model. Cell, 160, 1061–1071.

Wohleb, E.S. (2016) Frontiers | Neuron–Microglia Interactions in Mental Health Disorders: “For Better, and For Worse”. Frontiers in Immunology, 7.

Woolf, Z., Stevenson, T.J., Lee, K., Highet, B., Macapagal Foliaki, J., Ratiu, R., Rustenhoven, J., Correia, J., Schweder, P., Heppner, P., Weinert, M., Coppieters, N., Park, T., Montgomery, J., Smith, A.M., Dragunow, M., Woolf, Z., Stevenson, T.J., Lee, K., Highet, B., Macapagal Foliaki, J., Ratiu, R., Rustenhoven, J., Correia, J., Schweder, P., Heppner, P., Weinert, M., Coppieters, N., Park, T., Montgomery, J., Smith, A.M. & Dragunow, M. (2025) In vitro models of microglia: a comparative study. Scientific Reports 2025 15:1, 15.

Wu, J., Chen, X., Zhang, J., Wettschurack, K., Robinson, M., Li, W., Zhao, Y., Yoo, Y.-E., Deming, B.A., Shu, Y., Abeyaratna, A.D., Que, Z., Du, D., Tegtmeyer, M., Yuan, C., Skarnes, W.C., Zhang, Z.-Y., Rochet, J.-C., Wu, L.-J. & Yang, Y. (2026) Human microglia in brain assembloids display region-specific diversity and respond to hyperexcitable neurons carrying SCN2A mutation. Science Advances, 12.

Yuan, X., Puvogel, S., Rhijn, J.-R.v., Ciptasari, U., Esteve-Codina, A., Meijer, M., Rouschop, S., Hugte, E.J.H.v., Oudakker, A., Schoenmaker, C., Frega, M., Schubert, D., Franke, B. & Kasri, N.N. (2023) A human in vitro neuronal model for studying homeostatic plasticity at the network level. Stem Cell Reports, 18.

Zheng, H., Xu, W., Yu, F., Xu, J., Jiang, M., Ma, N., Shao, Z. & Ma, S. (2025) Protocol to generate species-specific astrocyte-conditioned medium for human organoid neuron maturation. STAR Protocols, 6.

Zhong, H., Xue, R., Han, Y., Liu, L., Zhao, J., Cai, M., Wang, S., Wei, P., Zhao, G. & Dong, H. (2025) S-ketamine exposure in early postnatal period induces social deficit mediated by excessive microglial synaptic pruning. Mol Psychiatry, 30, 3615–3631.

