## Supplemental Figures for "A human neuron-microglia tri-culture platform to study the influence of microglia on developing neuronal networks *in vitro*"

**Supplemental Information**

**Supplemental Figures 1 and2**

**Supplemental Table 1-6**

**Supplemental Video 1 - Legend**

### Supplementary Figures

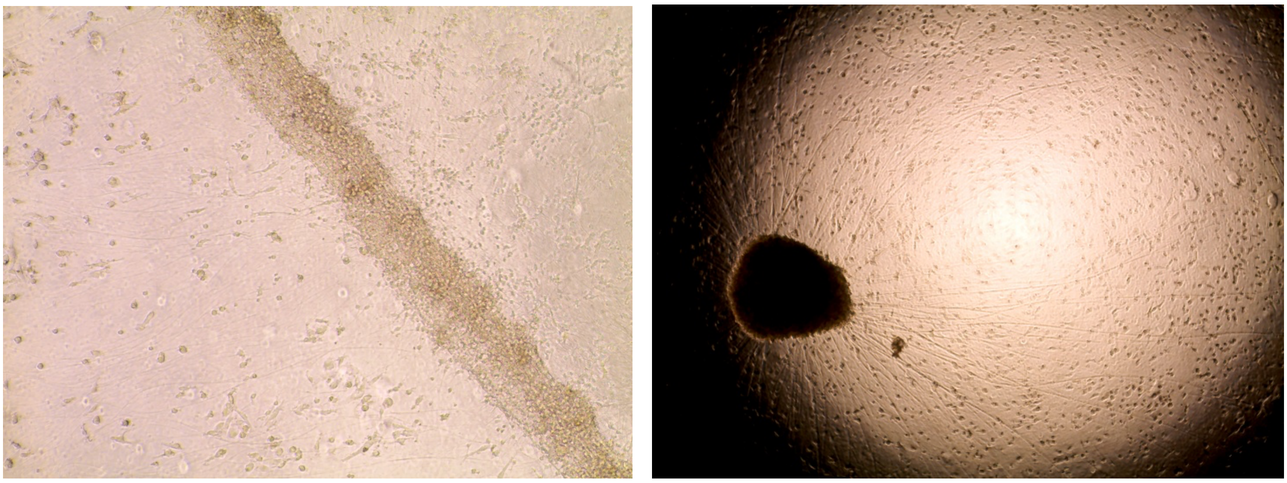

**Supplementary Figure 1.** Neural migration and clustering. Brightfield images showing migration and clustering problems occurred during optimisation of the protocol.

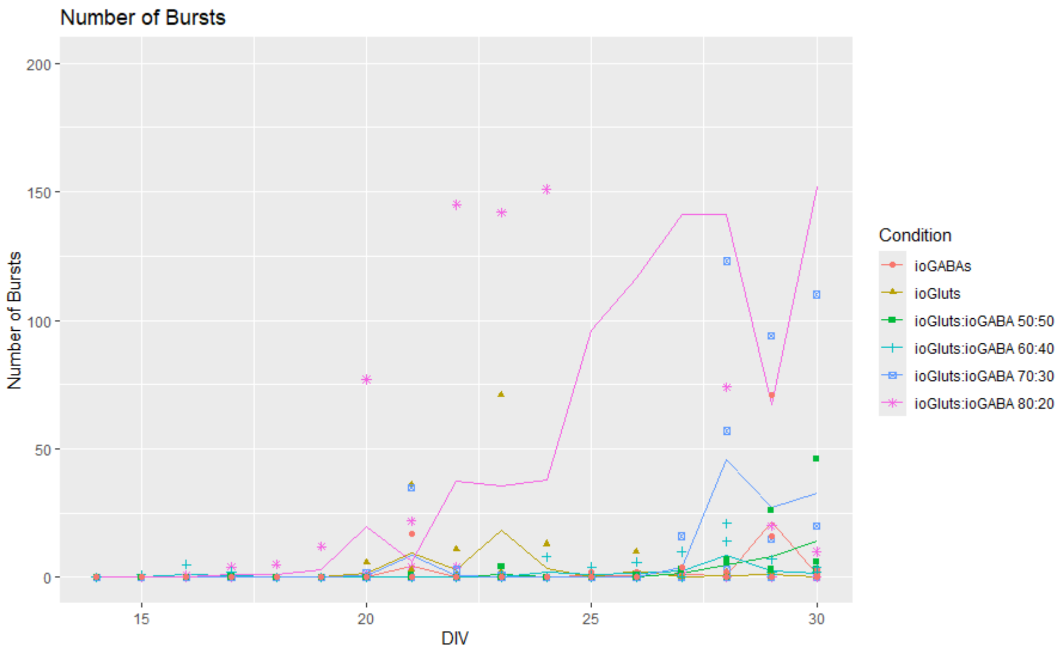

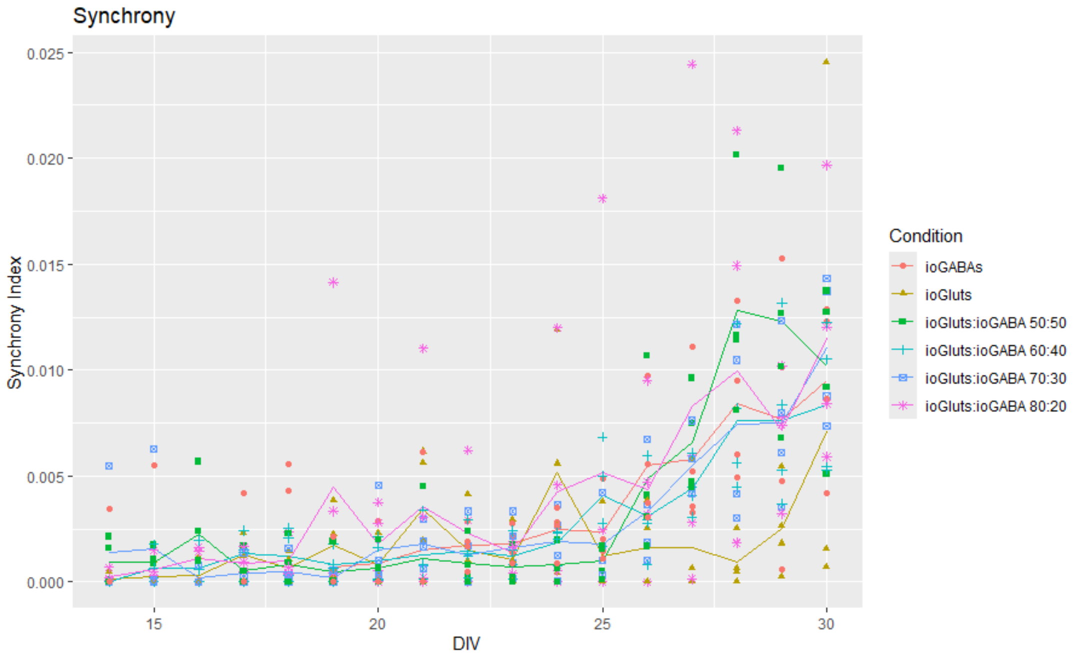


**Supplementary Figure 2.** Optimisation of cell ratio. Line graphs showing optimal number of bursts in the 80:20 condition, and comparable results in the Synchrony Index.

***Supplementary Table 1**. Antibodies used for immunocytochemistry*

| **Antibody** | **Company** | **Cat number** | **Dilution** |
|---|---|---|---|
| Anti-MAP2 antibody - Neuronal Marker | Abcam | ab32454 | 1:1000 |
| GABA Polyclonal Antibody | Invitrogen | PA5-32241 | 1:1000 |
| Anti-Iba1 Antibody | Antibodies.com | A82670 | 1:300 |
| Guinea pig Anti-NMDAR1 (GluN1) (extracellular) Antibody | Alomone Labs | AGC-001-GP | 1:1000 |

| | | | |
|---|---|---|---|
| Bassoon (D63B6) Rabbit mAb | Cell Signaling Technology | 6897S | 1:200 |
| VGAT Polyclonal Antibody | Invitrogen | PA5-27569 | 1:500 |
| Neuroligin2 antibody | Synaptic Systems | 129511 | 1:500 |
| Donkey anti-Chicken IgY (H+L) Highly Cross Adsorbed Secondary Antibody, Alexa Fluor™ 568 | Invitrogen | A78950 | 1:750 |
| Donkey anti-Goat IgG (H+L) Highly Cross-Adsorbed Secondary Antibody, Alexa Fluor™ Plus 647 | Invitrogen | A32849 | 1:750 |
| Donkey anti-Rabbit IgG (H+L) Highly Cross-Adsorbed Secondary Antibody, Alexa Fluor™ 488 | Invitrogen | A21206 | 1:750 |
| Goat anti-Rabbit IgG (H+L) Cross-Adsorbed Secondary Antibody, Alexa Fluor™ 568 | Invitrogen | A11011 | 1:750 |
| Goat Anti-Chicken IgY H&L (Alexa Fluor® 647) | Abcam | Ab150175 | 1:750 |
| Goat anti-Guinea Pig IgG (H+L) Highly Cross-Adsorbed Secondary Antibody, Alexa Fluor™ 488 | Invitrogen | A11073 | 1:750 |

| | | | |
|---|---|---|---|
| Goat anti-Mouse IgG (H+L) Cross-Adsorbed Secondary Antibody, Alexa Fluor™ 568 | Invitrogen | A11004 | 1:750 |
| Goat anti-Rabbit IgG (H+L) Cross-Adsorbed Secondary Antibody, Alexa Fluor™ 488 | ThermoFisher | A11008 | 1:750 |

**Supplemental Table 2** - Statistical analysis of imaging experiments from Figure 2.

**Supplemental Table 3** - Statistical analysis of imaging experiments from Figure 3.

**Supplemental Table 4** - Statistical analysis of MEA experiments from Figure 4.

**Supplemental Table 5** - Statistical analysis of imaging experiments from Figure 5.

**Supplemental Table 6** - Statistical analysis of MEA experiments from Figure 5.

**Supplementary Video 1. Calcium imaging of ioGlutamertgic neurons, ioGABAergic neurons and ioMicroglia cultured in Advanced DMEM/F12-based media.** Representative video of glutamatergic, GABAergic and microglia tri-cultures, grown in Advanced DMEM/F12-based media. Tri-cultures did not display spontaneous Ca2+ transients.
